# Structural Perturbations of Exon Skipping Edits within the Dystrophin D20:24 Region

**DOI:** 10.1101/2020.10.10.334763

**Authors:** Xin Niu, Nick Menhart

## Abstract

Exon skipping is a disease modifying therapy that operates at the RNA level. In this strategy, oligonucleotide analog drugs are used to specifically mask specific exons and prevent them from inclusion in the mature mRNA. Of course, this also results in loss of the corresponding region from the cognate protein, which is one possible therapeutic aim. Exon skipping can also be used to restore protein expression in cases where a genetic frameshift mutation has occurred, and this how it is applied to Duchenne muscular dystrophy, DMD. DMD most commonly arises as a result of large exonic deletions that juxtapose flanking exons of incompatible reading frame in the dystrophin gene, creating a frameshift and abolishing protein expression. Loss of dystrophin protein leads to the pathology of the disease, which is severe, causing death generally in the second or third decade of life. Here, the primary aim of exon skipping is the restoration of the reading frame by skipping an exon adjacent to the patient’s original defect. However, the therapeutically expressed protein is of course edited, and missing both the region of the underlying genetic defect, as well as the therapeutically skipped exon. While restoring some protein expression is good, how removing some region from the middle of a protein effects its structure and function is unclear. Complicating this in the case of DMD is the fact that the dystrophin gene is very large, containing 79 exons. Many different underlying deletions are known, and exon skipping can be applied in many ways. It has previously been shown that many exon-skip edits result in structural perturbations of varying degrees. What has been unclear is whether and how exon editing can be done to minimize these perturbations. In this study we examine a systematic and comprehensive panel of possible exon edits in a region of the dystrophin protein, and identify for the first time, exon edits that appear to maintain structural stability similar to wildtype protein. We also identify factors that appear to be correlated with the degree of structural perturbation, such as the number of cooperative protein domains, as well as how the underlying exon structure interacts with the protein domain structure.

## Introduction

Duchenne muscular dystrophy, DMD, is a genetic disease caused by the absence of the muscle stabilizing protein dystrophin. Being X-linked, it afflicts approximately 1 in 3500 in male births (1), and those afflicted have a dire prognosis of being wheelchair bound by their teenage years and have a median age of death of 21(2–4). Screening and other genetic techniques such as PIGD can have limited impact since it has a very high frequency of de novo mutation due to the extremely large size of the dystrophin gene size (2.5Mbp or 0.7% whole human genome)(5). The most common type of DMD causing gene defect is a large deletion (6) of one or more exons which results in juxtaposition of flanking exons with incompatible reading frame; this leads to a reading frame shift. A reading frame shift scrambles the amino acid sequence of the protein from that point on, and also inevitably in a protein as large as dystrophin creates new stop codons terminating the protein prematurely (5, 7).

However the very large size of the dystrophin gene responsible for its high incidence is put to advantage in the only current curative therapy approved for DMD, exon skipping (8–10). Because of the large size of the gene (~2.4 Mbp), deletions are generally small (1-100 kbp) in comparison, and much of the gene remains functional - but only if the reading frame can be restored, allowing the undamaged downstream C-terminal part of the protein to be expressed. Exon skipping essentially aims to restore the reading frame by skipping over an adjacent exon to restore the reading frame. This leads to a larger, but in-frame deletion (the patient’s original defect plus the skipped exon). This is not always possible, but where is it, is allows for some dystrophin protein expression, albeit in a modified form The first exon skipping drug, eteplirsen was approved in 2016 (11), with several others approved recently (12, 13) and other in late stage clinical trials.

Because this therapy results not in a full restoration of native dystrophin, but restoration of a modified dystrophin with a small deletion, to understand its potential we might examine natural occurrences of this situation. This is precisely the case Becker muscular dystrophy, BMD, where natural in-frame deletions (or more rarely other types of mutation) also produce modified dystrophin proteins (14, 15). BMD varies in severity from nearly as bad as DMD to nearly benign and it has long been suspected that this is related to the nature of the specific protein dystrophin defect arising from different genetic defects. However, due the large and complex nature of the DMD gene, which has 79 exons, there are a very large number of possible underlying exon deletions. While DMD and BMD are unfortunately common, in many cases individual defects are known only in a small number of patients. Furthermore, this condition has high variability, and even among patients with the same protein defect, and it has been challenging to understand which defects are “good” (less severe disease) and which “bad” (more severe disease)

For a few of the more common BMD exon deletions, sufficient case studies are available to show a definite relationship of the defect type to disease progression. For instance, in one study (16) examined three of the most common defects, with cohort sizes of 50, 38 and 18 patients, and was able to demonstrate a delay of slightly more than 10 years in the median age of onset of dilated cardiomyopathy between the “good” (deletion of exons 45 to 48, Δe45-48, age of 38.2) and the “bad” edit (Δe45-47, age of 27.6). However, this correlation is only seen on average over many patients; individual patients with these defect progress quite variably. For example, in that study fully 23% of patients in the “good” cohort had an age of onset below the median age for the “bad” edit (more severe progression); and 34% of patients in the “bad” cohort had an age of onset above the median age for the “good” edit (less severe progression). The reason for this variability is at present unclear, but may involve the interaction of specific defect with the individual genetic environment of the patient, for instance the interaction with many mi-RNAs impacting expression of dystrophin (17). However, it is clear that in some cases, the nature of the edit on dystrophin protein structure can have a significant clinical impact.

Whatever the reason, this high variability also means that for the majority of specific deletions, which are known only in a few patients, or with incomplete clinical data, it is difficult or impossible to make meaningful comparisons. To deal with this, other efforts have pooled defects into groups that are related by how the exon deletion interacts with the structure of dystrophin. Dystrophin is a large rod shaped protein composed of an N-terminal actin binding domain that connects to thin filaments in the sarcomere, a C-terminal globular region that binds to a large multiprotein complex embedded in the membrane; and in between a large central rod domain consisting of 24 tandem copies of a domain called a spectrin type repeat, STRs (18). Dystrophin then provides a mechanical connection from actin filaments to the sarcolemma, without which muscle tissue deteriorates over repeated contraction. Since the central rod domain makes up most of the gene and the protein, most mutations fall in this region and it is the impact of such edits on structure on this region that is the focus of this study. Although the rod region contains many binding sites for other proteins and biomolecules which are important for full functionality, it has been shown that this mechanical connection is the primary function of dystrophin, and disruption or the loss of this connection leads to muscle deterioration in DMD (19).

The 24 STRs that comprise this rod are themselves rod like domains, consisting of three bundled α-helices in a zigzag arrangement. Each STR roughly – but not exactly – corresponds to two exons (Figure S1). When exons are deleted in even numbers, a roughly whole number of STRs are removed but when in odd numbers, a fraction STR remains behind. The former situation, produced rod with intact STRs, and been termed *in-phase*. The latter situation, leaves a fraction domain at the defect site which is presumably disordered and unable to form an STR. This has been termed *out-of-phase* (in regard to the STR structure; not to be confused with in-frame and out-of-frame with regard to the codon reading frame). By grouping defects in this fashion, it has been possible to pool many different, rarer defects and achieve statistical power. *Out of phase* defects with an uncomplemented fractional STR region that might be expected (and are predicted by modelling techniques) to be disordered are associated with pooper clinical outcomes, especially more rapid progression to DCM by more than 10 years (20).

Within the more slowly progressing *in-phase* group, we may make a further distinction. Within STRs, the two exon boundaries occur either near the middle, or near the STR junctions. When removing an even number of exons, if the edit occurs at sites near the junction, this neatly excises the intervening STRs – essentially creating a shorter rod missing one or more STRs. However, when it occurs are middle exon boundaries, this fuses the N-terminus of one STR to the C-terminus of another STR, creating a non-native structure called a hybrid STR (21). The impact of this with regard to clinical outcome has not to our knowledge been studied. However, a number of biophysical studies on hybrid STRs have shown that while in certain cases, they can form stable STRs, they are generally of somewhat lower stability than native STRs ^7-10^. It has also been shown through biophysical and computational studies that the impact of an edit is can perturb the structure well away from the edit site, in particular at adjacent STR junction regions, even for hybrid STR with edit sites away from the junctions (22).

So, although it is clear that the nature of the structure at the edit site has some impact on the functionality of dystrophin and this is then linked to clinical outcome in some fashion, it the details of how any given edit will manifest are unclear, apart from a handful of cases. In order to examine this, we embarked on a comprehensive scan of a region of the dystrophin rod, producing all possible exon edits in a systematic way. We chose the D20:24 domain within the rod region, which is in the latter half of hotspot 2 (HS2) region, the larger of the two hotspot regions where most patient defects are found. This region is thought to be a relatively independent section of the rod, bounded on one site by the H3 region and on the other by the C-terminal globular region, and makes a convenient yet feasible region to do a comprehensive scan. Overall, there are 63 possible frame conserving edits in HS2, and in the D20:24 region there are 11 (Figure 1), making a full scan more feasible. In this set there also 4 sets of alternative repairs (different repairs that may be produced by single exon skipping of the same underlying defect with AONs targeting different exons) what are especially clinically relevant. In these cases, for patients with these underlying defects, exon skipping may be applied to produce different, mutually exclusive repairs. Understanding which repair might be superior is crucial as additional exon skipping reagents are developed.

**Figure 1.**
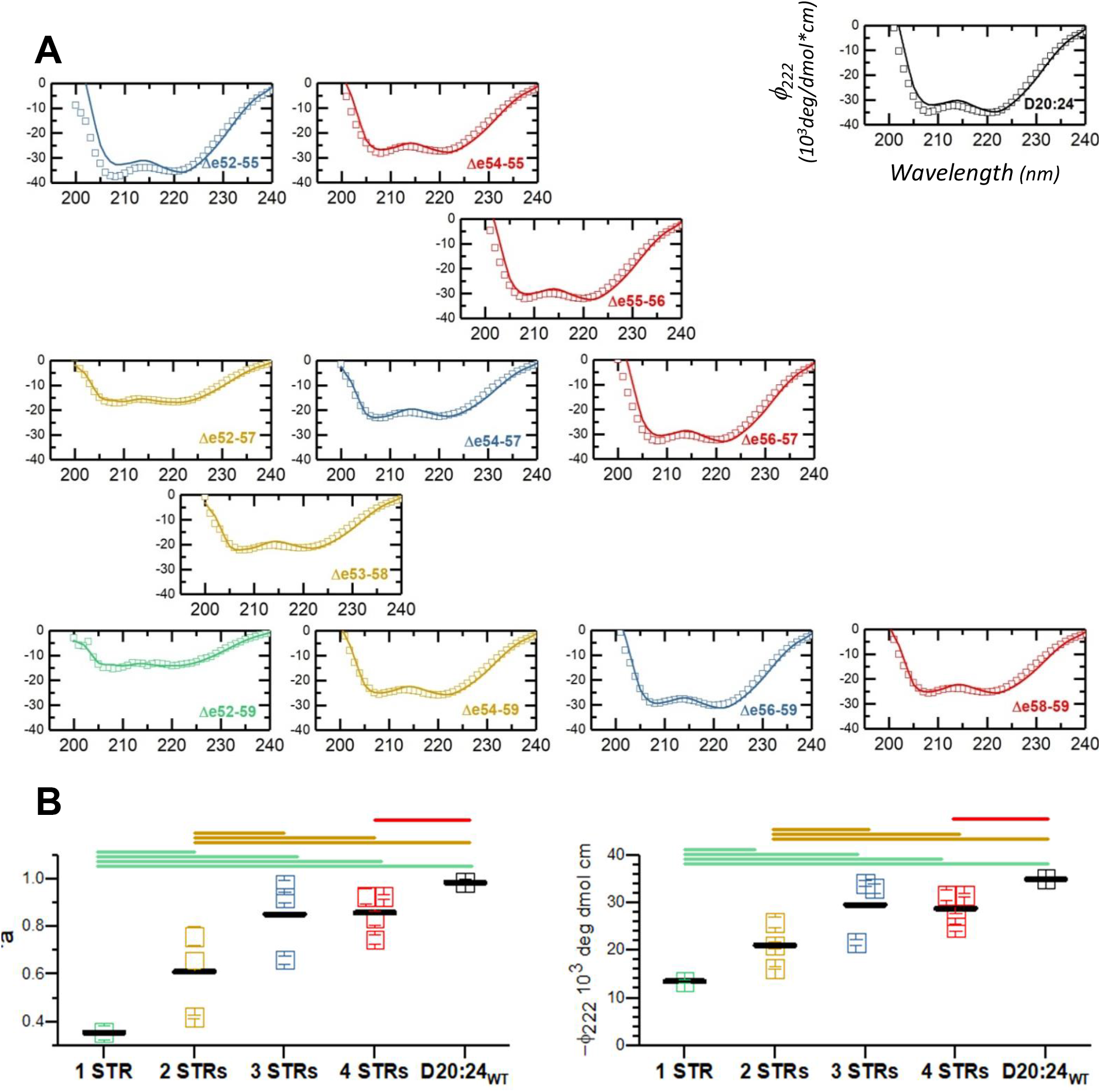
Helicity and α-helical content. **A** CD spectra of target proteins are shown, and presented laid out by first exon skipped, 52 to 58, along the horizontal coordinate; and last exon skipped, 55 to 59 along the vertical coordinate. This results in target proteins of varying sizes containing 1 to 4 STRs (see figure S1) diagonally, which are presented in different colors: 1-STR green, 2-STR gold, 3-STR blue, 4-STR red. Wild type unskipped D20:24 is in black at upper right. These spectra are fit as described (solid line) to produce a fractional helical content, f_α_. **B** We present both f_α_ and φ222 as a function of STR number, as well a size cohort means. Statistical significance (P< 0.005) is shown by bars above.

Beyond this very practical comparison, this comprehensive scan allows us to examine the fusion of certain exons with several other exon partners at multiple edit sites, and begin to understand which edits are viable, and why certain edits are stable and certain destabilizing. Since structural factors linked to exon edit type have been linked to clinical outcome, it is hoped that a better understanding of how to develop well-edited dystrophins will lead to improved therapies, both for exon editing, as well as other therapies involving edited dystrophins, such as mini-dystrophin gene therapy, or CRISPR mediated gene editing.

## Experimental Procedure

### Cloning and Protein production

Proteins were cloned, expressed and purified as previously described (23). Briefly, this involved expression in *E. coli* from a pGEX plasmid-based expression system, with target deletions constructed by PCR of appropriate fragments and Gibson assembly. The exact details of the end points of each protein constructed were given in Table 1 and Figure S2. Proteins were isolated by a double-tag affinity chromatography protocol using an initial glutathione-S-transferase, GST, affinity step, followed by removal of the large GST fusion tag by thrombin cleavage, and then a His-tag cobalt-immobilized metal affinity step. In the absence of any degradation during this initial extraction, only full-size fragments would be produced by this double tag process.

**Table 1.** Details of targets constructed in the D20:24 region. All targets start using the same N- and C-termini as previously used for wild type rod studies, immediately before D20 and after D24. The edit sites result in an edit site sequence as shown, with pink residues being from the N-terminal side and blue residues being from the C-terminal side; these are the same colors used in the coloring scheme for Figure S1 where Robetta models of these edits are shown. The vertical bar | shows the edit junction, except in the cases of Δe53-58 and Δe55-56 where this occurs in the middle of a codon, which created a new amino acid belonging to neither the N- nor C-termini shown in purple.

| Edits Name | STR # | N-terminus | Edit Site | C-terminus |
| --- | --- | --- | --- | --- |
| D20:24 | 5 | SSLML | N/A | DFGPA |
| Δe54-55 | 4 |  | TETK \| DLQG |  |
| Δe55-56 | 4 |  | SIHKRSHLE |  |
| Δe56-57 | 4 |  | KQWQ \| AFKR |  |
| Δe58-59 | 4 |  | DVHR \| ALRG |  |
| Δe52-55 | 3 |  | IKQK \| DLQG |  |
| Δe54-57 | 3 |  | TETK \| AFKR |  |
| Δe56-59 | 3 |  | KQWQ \| ALRG |  |
| Δe52-57 | 2 |  | IKQK \| AFKR |  |
| Δe53-58 | 2 |  | ITDRKLPPE |  |
| Δe54-59 | 2 |  | TETK \| ALRG |  |
| Δe52-59 | 1 |  | IKQK \| ALRG |  |

However, in some cases, especially with less stable targets, some degradation was seen, presumably by endogenous proteases from the *E. coli* expression platform. As such, when needed, purification was continued by ion exchange and hydrophobic interaction exchange chromatography, as shown in Table 2. In the ion exchange step, proteins were loaded onto a Hi-Trap Q (GE Lifesciences) column in 25mM of tris pH 8.5 and eluted with a gradient of NaCl, with elution [NaCl] concentration as shown in Table 2. For hydrophobic interaction chromatography samples were loaded onto a Hi-Trap Octyl FF column (GE Lifesciences) in 25mM phosphate buffer at pH 6.3 with 1M NH4SO4, and eluted with a gradient to 25mM phosphate buffer at pH 6.3 with 30% acetonitrile. The percentage acetonitrile of elution is similarly given in Table 2. Final protein purity was always >95% as assessed by SDS-PAGE.

**Table 2.** Target specific purification conditions for ion-exchange and hydrophobic interaction chromatography

| Edits Name | IEC Salt Concentration | HIC Elution CH<sub>3</sub>CN % |
| --- | --- | --- |
| D20:24 | 300mM | 9% |
| Δe54-55 | N/A | 4% |
| Δe55-56 | 200mM | N/A |
| Δe56-57 | 225mM | N/A |
| Δe58-59 | 175mM | N/A |
| Δe52-55 | 200mM | N/A |
| Δe54-57 | 250mM | N/A |
| Δe56-59 | 150mM | N/A |
| Δe52-57 | 175mM | N/A |
| Δe53-58 | 225mM | N/A |
| Δe54-59 | N/A | N/A |
| Δe52-59 | N/A | N/A |

### Computational Model Construction

The primary sequences of the target edits, and the D20:24 wild type parent molecule (see Table 1) were submitted to the Robetta automated protein modelling server (24).

### Biophysical Characterization

#### Helicity

Total helicity was assessed by circular dichroism (CD) as previously described (23). The CD instrument was calibrated against camphor sulfonate primary standards according to the manufacturer’s instructions before all the experiments. Spectra were acquired using a Jasco J-710 spectrometer at a protein A_280_ ≅ 0.5 in PBS buffer (5mM sodium phosphate at pH 7.4 plus 150mM NaCl) in a 1.0 mm path cell from 190 nm to 300 nm. The resulting spectra converted into molar ellipticity, φ_222_, using a molar extinction coefficient provided by Protparam and fit to a 3-basis model (25) that we and others (26–28) have used extensively for assessing STR containing proteins.

#### Thermal denaturation

Thermal denaturation experiments were conducted to assess thermodynamic stability. A protein sample with ellipticity of ~50 mdeg (A_280_ ~ 0.5) with was heated from 0.5°C to 98.5°C while monitoring the CD signal as 222 nm as previously described (29). This data was processed to the derivative of the molar ellipticity with respect to temperature, dφ_222_/dT and fit to a two state unfolding (30) to provide melting temperature, Tm, and enthalpy of unfolding, ΔH.

#### Protease challenge

Sensitivity to proteolysis was performed as described (31). This is correlated with the presence of disordered regions, and we and others have used this to assess a wide range of in native and edited STR rods. Briefly, 2.5 ug protein samples were incubated with the non-specific protease Proteinase K (PK) at various geometrically decreasing concentrations in PBS for 30 min at 37C. Proteolysis was stopped by denaturation at 99 C for 5 minutes, and samples and were then analyzed by SDS PAGE. Undigested full-length protein was quantified by densitometry and the fraction remaining, *f*, was calculated and fit to a first order decay process *f* = e^−[PK]/PK_50_^. The result was reported as PK_50_ value.

#### Hydrodynamic radius by dynamic light scattering

Dynamic light scattering using a Malvern Zetasizer was used to measure the hydrodynamic radius of the targets to assess rod like shape and aggregation. Hydrodynamic radius was compared previously expressed unedited STR proteins, as well as globular protein standards (lysozyme, 14 kDa, ovalbumin, 45 kDa, and yeast alcohol dehydrogenase, 114 kDa) as previously described (29). Hydrodynamic radius scales with MW^1/3^, but is larger for rod-shaped proteins than globular proteins (32, 33).

#### Data Reproducibility and Significance Testing

The standard of date reproducibility was that a data of all the tests were acquired from at least three independent experiments using at least two independent batches of protein. Comparisons for statistical significance are presented at P<0.05 (significant) or P<0.005 (highly significant) as indicated, using a two tailed Student’s t-test, as well as effect size as measures by Cohen’s d parameter, d> 2.

## Results

### Secondary Structure by CD

All proteins showed circular dichroism (CD) spectra with two peaks at ~222 nm and ~208 nm, as shown in Figure 1, which is consistent with a highly α-helical structure expected of STR motifs. These spectra were analyzed for a-helical content by a 3-basis sets method (34) used by us and others (27, 28, 35) for STR proteins, which is presented in Table 3.

**Table 3.** Summary of parameters measured for all targets. Data is presented as mean +/- standard error.

| target | STR # | class | $f_a$ | $\phi_{222}$<br>mdeg/μmol cm | $\phi_{222}/208$ | $T_m$<br>°C | DH<br>kJ/mol | PK50<br>ng | $r_H$<br>nm |
| --- | --- | --- | --- | --- | --- | --- | --- | --- | --- |
| D20:24 | 5 | native | 0.98 ± 0.01 | -35.0 ± 0.3 | 0.996 ± 0.004 | 70.0 ± 0.2 | 467 ± 18 | 3.42 ± 0.18 | 8.67 ± 0.17 |
| Δe54-55 | 4 | hybrid | 0.83 ± 0.03 | -27.2 ± 0.5 | 0.963 ± 0.007 | 65.3 ± 0.5 | 261 ± 11 | 0.13 ± 0.01 | 8.03 ± 0.35 |
| Δe55-56 | 4 | junction | 0.93 ± 0.03 | -31.6 ± 1.2 | 0.970 ± 0.005 | 64.6 ± 0.2 | 889 ± 72 | 7.40 ± 0.14 | 7.53 ± 0.63 |
| Δe56-57 | 4 | hybrid | 0.93 ± 0.01 | -31.6 ± 0.4 | 0.962 ± 0.006 | 71.0 ± 0.1 | 398 ± 36 | 0.21 ± 0.04 | 7.37 ± 0.21 |
| Δe58-59 | 4 | hybrid | 0.75 ± 0.02 | -24.8 ± 0.7 | 0.963 ± 0.003 | 71.1 ± 0.2 | 495 ± 68 | 0.64 ± 0.03 | 9.05 ± 0.26 |
| Δe52-55 | 3 | hybrid | 0.98 ± 0.02 | -34.1 ± 0.7 | 0.927 ± 0.004 | 65.8 ± 0.8 | 269 ± 11 | 0.27 ± ##### | 8.17 ± 0.60 |
| Δe54-57 | 3 | hybrid | 0.66 ± 0.02 | -21.6 ± 0.6 | 0.928 ± 0.019 | 65.1 ± 0.2 | 183 ± 11 | 0.07 ± 0.01 | 7.27 ± 0.17 |
| Δe56-59 | 3 | hybrid | 0.92 ± 0.02 | -33.0 ± 0.9 | 1.014 ± 0.006 | 71.8 ± 0.4 | 495 ± 23 | 5.14 ± 0.61 | 6.69 ± 0.13 |
| Δe52-57 | 2 | hybrid | 0.42 ± 0.01 | -16.2 ± 0.2 | 0.957 ± 0.003 | 64.9 ± 1.5<br>42.3 ± 2.8 | 165 ± 29<br>74 ± 5 | 0.43 ± 0.10 | 7.03 ± 0.61 |
| Δe53-58 | 2 | hybrid | 0.66 ± 0.03 | -21.0 ± 0.7 | 0.903 ± 0.029 | 65.0 ± 0.6 | 171 ± 15 | 0.66 ± 0.01 | 7.61 ± 0.41 |
| Δe54-59 | 2 | hybrid | 0.76 ± 0.04 | -25.9 ± 1.1 | 0.987 ± 0.005 | 64.1 ± 0.3<br>33.9 ± 0.5 | 150 ± 7<br>96 ± 7 | 0.29 ± 0.01 | 5.57 ± 0.07 |
| Δe52-59 | 1 | hybrid | 0.36 ± 0.03 | -13.5 ± 0.3 | 0.860 ± 0.035 | 45.6 ± 1.8 | 54 ± 5 | 0.36 ± 0.07 | 13.33 ± 1.37 |

For the parent molecule an ellipticity φ_222_ = −35.0 mdeg/dmol cm^2^ was observed, corresponding to 98% helicity when This is consistent with the highly alpha helical structures expected of STR rods, and is in fact perhaps too high. With two non-α-helical turns for each STR, we might expect >10 residues out of the ~117 in each STR to be non-helical, for an upper bound of ~92% helical. However, a number of factors make secondary structure by CD somewhat uncertain. For instance, supercoiling – present in all knowns high resolution structures of STRs - is known to distort the typical a-helical spectra, increasing signal at the 222nm peak (36) and resulting in overestimated helical content relative to methods based on references sets with mostly linear and shorter helices. Comparison of many methods, including more sophisticated algorithm suggests an overall uncertainty of at best 10% when used on proteins where X-ray structures are known (37); or see for instance figure 4 in (38).

**Figure 2.**
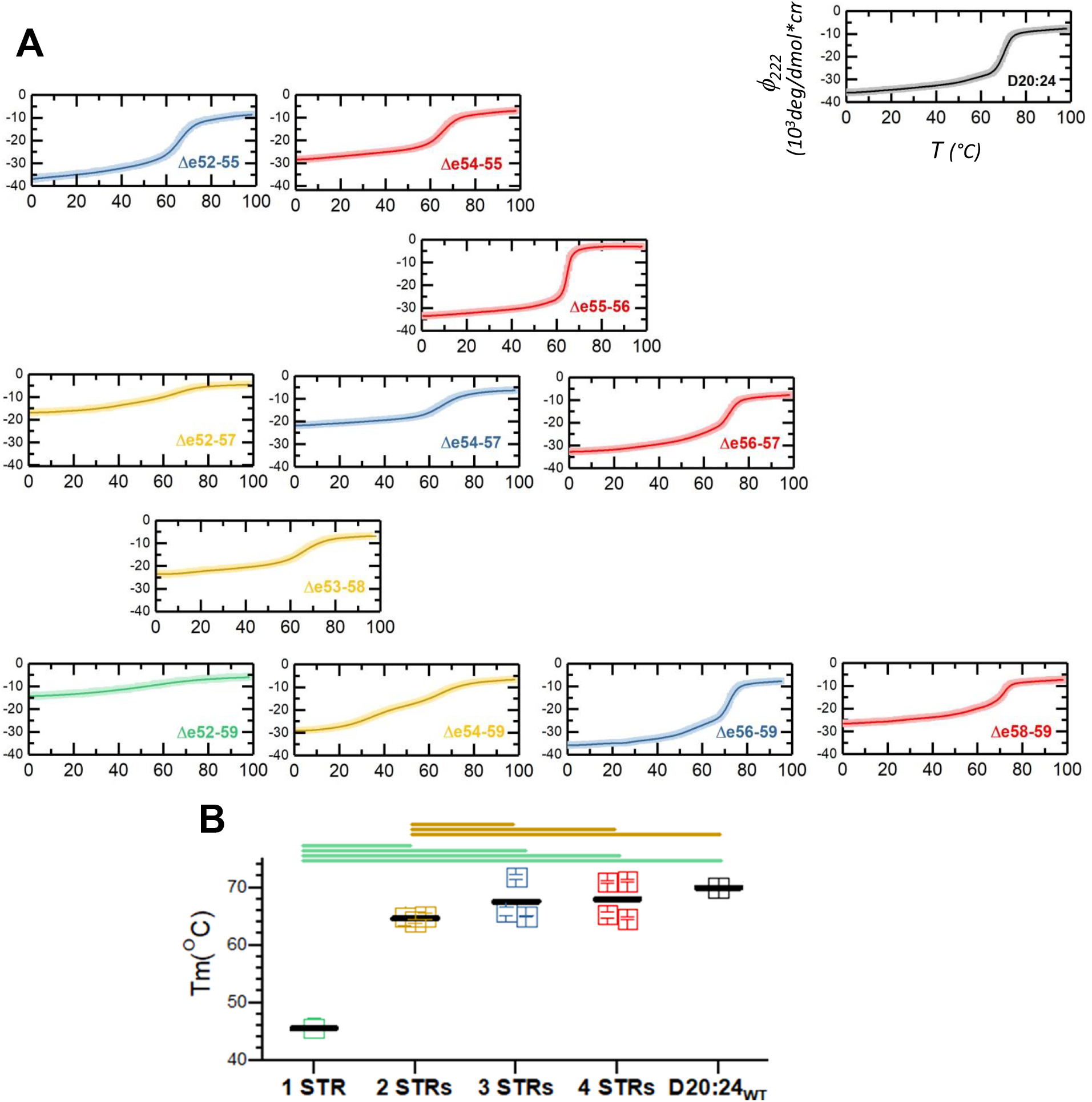
Thermal stability. **A** Proteins were subject to thermal unfolding and monitored by CD signal and 222nm. This yielded unfolding curves that were FFT filtered, solid lines, and then subject to thermodynamic analysis (Figure 3). **B** Tm as determined by thermodynamic analysis presented as a function of STR number, with cohort means also presented (black lines). Statistically significant differences (P<0.05) between cohorts is shown by bars above.

**Figure 3.**
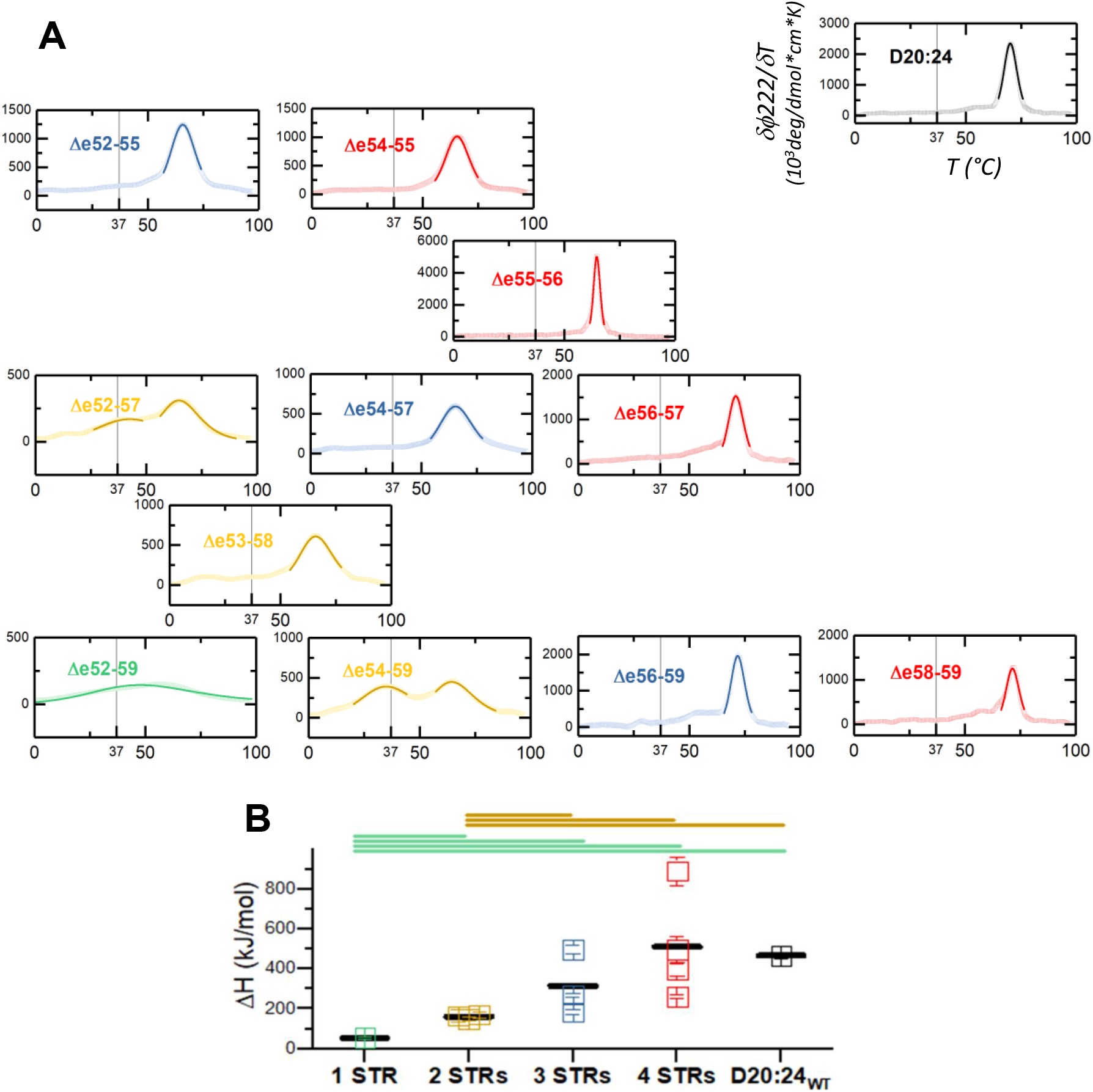
Thermodynamic analysis of thermal stability. **A** Unfolding curves (Figure 2) were converted to derivative form δφ_222_/δT, and then fit to a two-state unfolding equations as described. In two cases, Δe52-57 and Δe54-59 two unfolding events were notes and separately fit. This yielded Tm and enthalpy of unfolding, ΔH, which is associated with transition width in the raw data and peak width here in the derivative data. **B** ΔH is presented as a function of STR number, with cohort means also presented (black lines). Statistically significant differences (P<0.05) between cohorts is shown by bars above.

**Figure 4.**
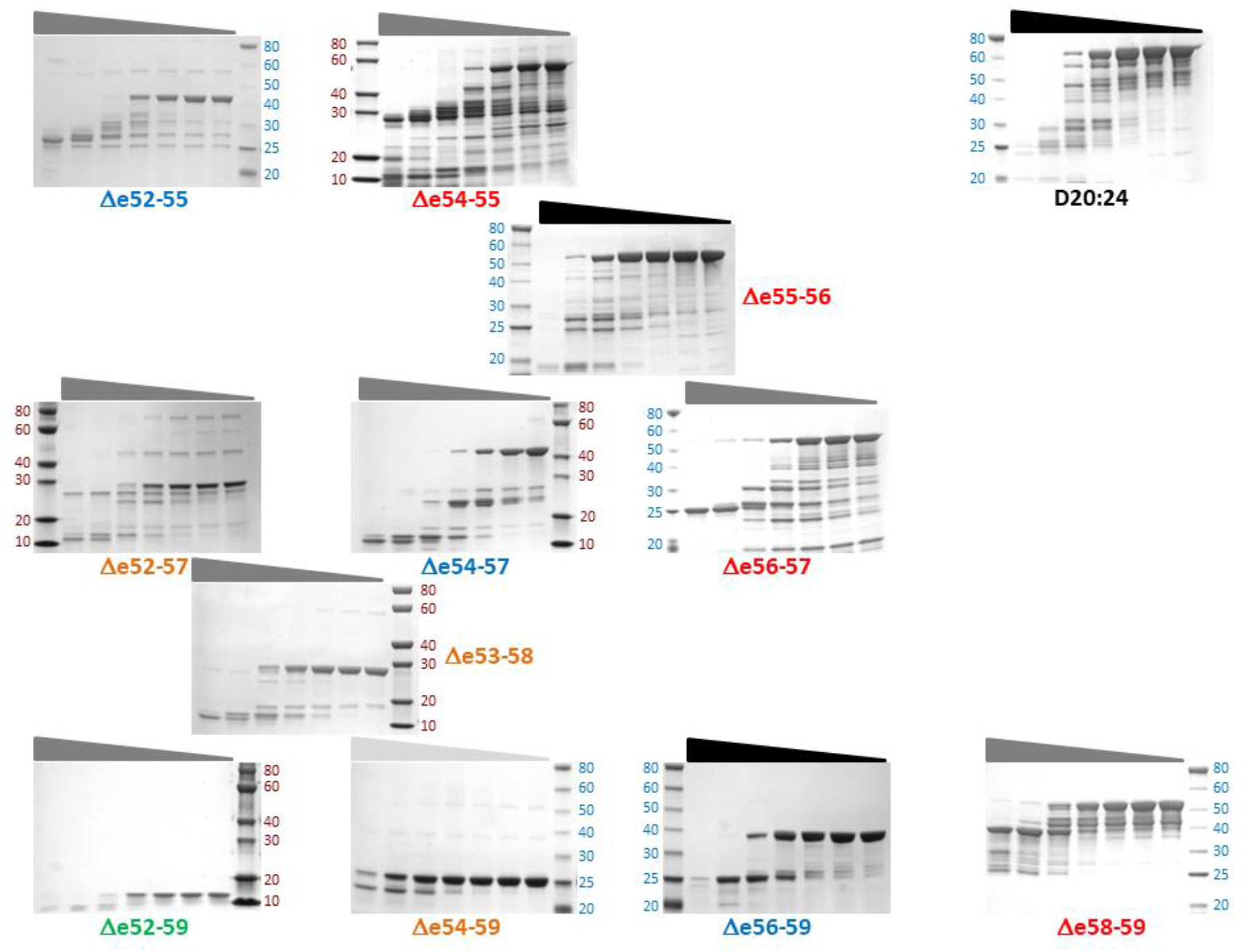
Protease challenge. Target rod proteins are subjected to various concentrations of proteinase K and analyzed by SDS-PAGE. Molecular mass standards are labeled as leftmost or rightmost lane. The grey/black triangle above the gel graph indicate different concentration of proteinase K varies from high to low from left to right, respectively, as shown in the graphs.

Here we use CD to examine differences between edited versions of the same protein. As such, idiosyncratic perturbations (i.e. caused by supercoiling, are expected to apply in similar fashion to all. What is more important are the differences seen between the various edits? Here we find that large differences are seen, with helicities ranging from a low of 36% (φ_222_ = −13.5 mdeg/dmol cm^2^) in the case of the smallest protein, the 1-STR Δe52-59, to a 98% value identical to wildtype in Δe52-55. In general, larger edits / smaller proteins exhibit lower helicities, consistent with less STR-like structure. Indeed, the second lowers helicity observes was one of the two STR proteins, Δe52-57 at 42%. All other edits, the 2 remaining 2-STRs variants, and all 3-STRs variants exhibited helicities > 66%. Groups into STR size cohorts, Figure 1, we see a monotonic increase in average helicity from the single example of 36% for the only one 1-STR variant (Δe52-59), to 63%, 85% and 86% for the averages of 2-3- and 4-STR edits. This may have implications for design of modified dystrophin rods, either by exon skipping or by other strategies that employ engineered dystrophin rod such as mini-dystrophin gene therapies.

However, another lesson presented by this regionally comprehensive scan is that such general conclusions need to be tempered by examination of the specific properties of any give edit. We see that, especially at the lower end, variation within cohorts can be more significant than this general trend. For instance, within the 2-STR cohort, we find as expected the comparatively low helicity Δe52-57 at 42%, but also Δe54-59 much higher at 76%. Among 3-STR rods, we see a near-wild-type 98% for Δe52-55, Δe56-59 at 92 % - but also Δe54-57 at only 66%. The four STR cohorts have less variation and a higher minimum value, varying between 75% and 93%.

### Protein Stability by Thermal Denaturation

We also assessed the thermodynamics of the protein structure by thermal denaturation, which provided the melting temperature, Tm, and enthalpy of unfolding, ΔH. Here once again we can use the unedited rod as a benchmark, which shows a single cooperative transition with Tm=70.0°C and an enthalpy ΔH=467 kJ/mol. For the edits, in most cases we also saw a single cooperative transition, but in two cases, Δe52-57 and especially, Δe54-59 we see not just one cooperative transition, but two roughly equal two transition suggesting that the edit split the rod into two independently unfolding regions that do not structurally cooperate.

We found that edits could be destabilizing, neutral, or in two cases. By Tm, seven were destabilized with Tm in the range 64°C to 66°C (Figure 2), and a further one, Δe52-59 was even further destabilized, with Tm=45°C. All those in the 64-66 range showed destabilization with respect to ΔH, falling in the range 150 kJ/mol to 261 kJ/mol (compared to 467 for wild-type D20:24); and Δe52-59 showed a very low ΔH transition of only 54 kJ/mol. However, one of these, Δe55-56 was significantly stabilized in terms of folding enthalpy, with ΔH=889 kJ/mol. A further 3 were marginally stabilized with Tm in the range 71°C to 72°C; and has ΔH’s in the near the wild type value, in the range 398 kJ/mol to 495 kJ/mol. When examined in the STR size cohorts, we fine the same trend as was observed for helical content. Smaller edits producing larger rods produced, in general, less perturbation in both Tm and ΔH. Once again, there are significant variations within size cohorts, especially in the 3-STR cohort.

Finally, we note that for some STRs, the combination of low Tm and low ΔH (as well as, in two cases, multiple transitions), means that some of these targets begin to unfold at the physiologically significant temperature of 37°C, indicated by the vertical lines in Figure 3. This includes the very unstable Δe52-59, as well as Δe52-57 and Δe54-59.

### Protein Stability and Folding by Protease K Challenge

Protein structure stability was also assessed by proteinase K (PK) challenge, which yields a PK_50_ value at which 50% degradation occurs in the standard assay, Figure 4. Unlike thermal denaturation which detects structured regions, protease sensitivity is correlated with disordered backbone regions, and so PK_50_ reports on unstructured regions that might be produced by the edits. As a baseline, the wild type D20:24 exhibited a PK_50_=3.4 ng. All but two of the exon edits exhibited lower PK values, ranging from 0.07 to 0.6 ng (Figure 5). On the other hand, two edits were more resistant: Δe55-56 at PK_50_=7.4 ng, and Δe56-59 at PK_50_ = 5.1 ng. These two edits are also stabilized as assessed by ΔH. We note that in the D2024 region, other, 2-STR wild-type protein have been studied and have higher PK values (D20:21 PK_50_ = 17ng; D21:22, 22 ng; D23-24 = 8 ng (23)). This may suggest that these edits simply remove a naturally occurring a marginally disordered region from the wild type rod. Finally, examining these in the STR size cohorts shows the same tread as observed in helicity thermodynamics: larger edits, equivalent to smaller rods, are more strongly perturbed. In this case, the heterogeneity in 3- and 4-STR cohorts is greater than the other techniques and we were not able to demonstrate a statistically difference between 1-, 2-, 3-, and4_STR cohorts. However, 1-STR and 2_STR targets were significantly more sensitive than the unperturbed D20:24 wild type rod (Figure 5B)

**Figure 5.**
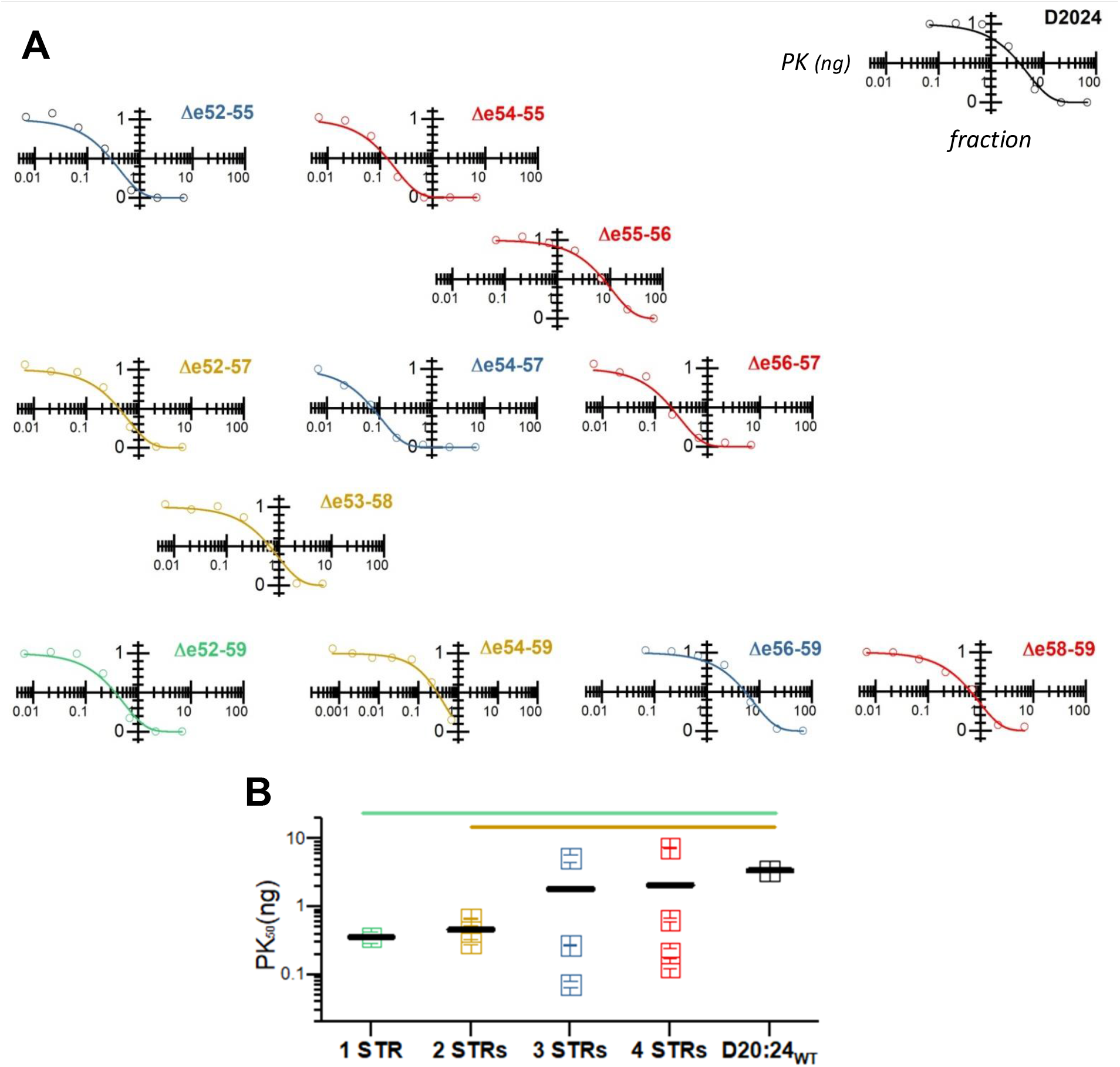
Protease sensitivity assay. **A** Target proteins were subject to varying amounts of proteinase K under a standard condition, and then analyzed by SDS PAGE. The amount of full-length protein remaining is assessed by densitometry and fitted as described to an exponential decay, yielding a half-life value, PK_50_. Only two edits, Δe55-56 and Δe56-59 exhibited higher PK_50_ values than D20:24 native rod. **B** The 1- and 2-STR proteins shows statistically significant enhanced sensitivity (P<0.005) as shown by the comparison bars above the graph.

### Protein Diameter Measured by Dynamic Light Scattering (DLS)

Hydrodynamic diameter of the target protein was measured by DLS (Table 3). As rods shaped proteins, STRs are expected to exhibit diffusional rates somewhat slower than globular proteins of similar molecular mass (39); this was confirmed in Figure 6, where the hydrodynamic radii are displaced from a standard curve created by globular proteins by 0.2 to 0.3 on a log scale (i.e. 70% to 100% larger). This is a similar offset observed for many other well-formed type 1- and 2-STR fragments of dystrophin, as well as some other exon edited STR rods. The sole exception to this was Δe52-59, which displayed an anomalously large hydrodynamic ratio outside of this range, at 13.2 nm. This very large rH suggest that this protein may aggregate in solution, something that has been long been observed for some STR rods (35), especially other edited rod, and even some less stable 1- and 2-STR rods when expressed independently. In conjunction with the low stability of Δe52-59 by other measure (thermal stability, proteolytic susceptibility) this further suggests this edit is strongly destabilizing.

**Figure 6.**
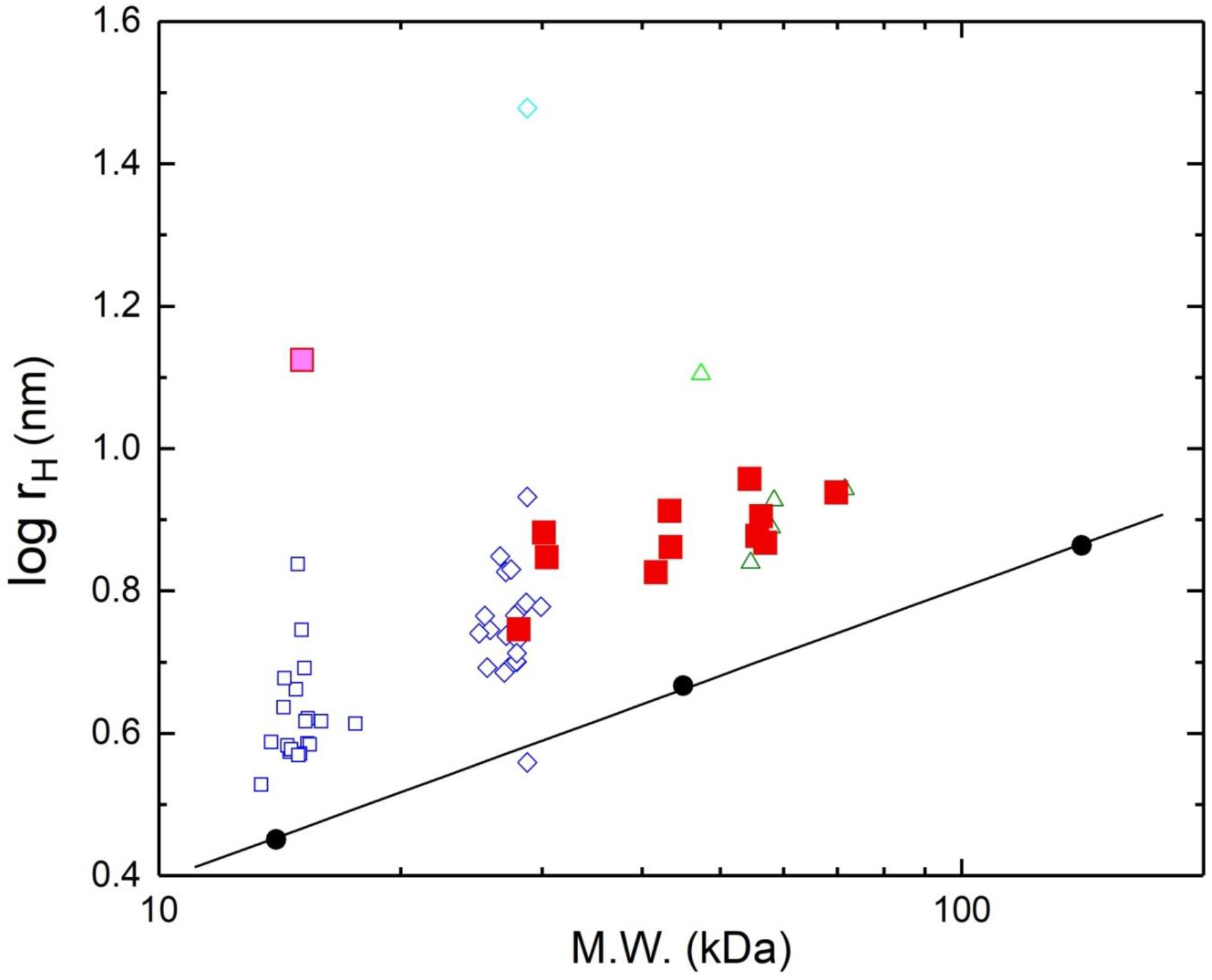
Hydrodynamic radius. Diameter of all constructs were measured by DLS and presented as a function of molecular mass. The 11 exon-edit targets and wildtype D20:24 are plotted as red solid squares, except for Δe52-29 which is shown as a red square with a magenta interior. For comparison, also shown is data from previous studies including native 1-STR (open squares) and 2-STRs (open diamonds) proteins, as well as other exon edited rods in the hinge 3 region (green open triangle). Three globular proteins are labeled in black solid circles (lysozyme; 14.4 kDa, ovalbumin 45 kDa; alcohol dehydrogenase, 114 kDa). Most STR proteins have r_H_ values that offset from globular proteins by similar amount, but a few show anomalously high r_H_ – in this study, Δe52-59.

## Discussion

One aim of this study was to determine which of all possible edits are “good” edits and which are “bad” edits. This is decidedly imprecise and unscientific phrasing, and we must of course explain what we mean by “good” vs “bad”. Ultimately, as edits proposed for DMD therapy we probably mean “therapeutically successful” which edits that restore as much function to a patient as possible. This holy grail is a challenging target, and ultimately requires assessment in a clinical setting. Clinical studies entail significant cost and more importantly, risk. To mitigate this, they are often informed by preclinical studies using surrogate markers to predict likely success, and to manage risk. One such surrogate marker is the impact of the edit on the protein’s structure. Here, multiple techniques have been used to assess this impact, and compare such edited protein to the native protein. We are primarily concerned with protein stability, but also secondary structure, since dystrophin is a largely α-helical protein with a simple secondary structure. We propose that those edits that most closely resemble undamaged, unedited dystrophin are likely to be most successful as clinical targets. This is supported by retrospective BMD studies that suggest disrupted protein structure is associated with worse clinical progressions (16, 20, 40, 41)

In this study, we focused on one region, the D20 to D24 regions delimited by H3 and H4. In this 5-STRs region, we studied 11 exon edits in a systematic fashion to determine if, and how, the nature of the edit impacted the proteins stability and structure; a summary of these results has already been presented in Table 2. Here, we seek to understand what we have learned from this. The first question is, “*Does the nature of the edit impact the proteins stability and structure?*” which we can answer with an emphatic yes.

In most cases, these rod edits behave very differently, and these differences are consistently manifest in studies that probe tertiary structure (thermal stability) secondary structure (fraction helical by CD) and disordered structure (proteolytic sensitivity) in all measures of stability. Of the 66 possible pairwise comparisons between the 12 studies, 50% showed statistical differences at P<0.05) in all four measured properties (fa, ΔH, Tm, PK_50_), and 85 % in at least three (Figure S3). For statistically significant differences (P<0.005), 18 % of the comparison showed variation in all four measurement techniques and 65% in at least three methods.

This shows that the edit impacts structure – but for better or worse? In terms of identifying more stable and less stable edits, two of them, Δe55-56 and Δe56-59 are the most stable and most comparable to native D20:24. Both of them are highly helical (Δe55-56: 93%, Δe56-59: 92%, D20:24: 98%) and show good thermodynamic stability (ΔH_unfolding_ for Δe55-56: 889kJ/mol, Δe56-59: 495kJ/mol, D20:24: 467kJ/mol) and resistance of proteolytic degradation (PK_50_ for Δe55-56: 7.40ng, Δe56-59: 5.14ng, D20:24: 3.42ng). On the opposite side, Δe52-29 is the most destabilized with the lowest helical content (36%) and thermodynamic stability parameters (Tm: 45.6°C, ΔH: 54kJ/mol). The other edits fall in between.

So, have we learned anything new? After all, we (21, 22, 29, 31) and others differences (16) have previously studied exon edited proteins before and shown stability. However, in all those cases the exon edits were significantly destabilized compared to wild type unskipped dystrophin. This is the first case, to our knowledge, that an exon edited protein exhibits stability all metrics testes comparable to, and even slightly above, that of dystrophin. This illustrated that fully functional deletions at can be achieved, at least from a protein structural perspective. The fact that 9 of 11 were somewhat, to significantly destabilized, does however confirm these previous findings that many exon edits, while potentially viable, are generally significantly disrupted.

A secondary aim of this work is to move beyond simply determining which edits are most and least perturbed, but also to identify any patterns or rationale for what makes a specific edit more or less similar perturbing. Above, we identified two edits that are superior in stability than the other edits, and similar to undamaged dystrophin. But why did this happen? What distinguished these particular edits?

### Helicity and Thermal Stability are correlated with STR number

One of the most obvious patterns that emerged is that all structural and stability metrics, f_α_, Tm, ΔH, and PK_50_, correlated with size, as shown in the size-grouped analysis of previous results (Figure 1B, 2B, 3B and 5B). Along with the 5-STR D20:24 unedited parent molecule, we studied four 4-STR targets, three 3-STR targets, three 2-STR targets, and only a single 1-STR, Δe52-59. Of course, there is some variation, and not all motifs in each size cohort exhibited identical properties, however a striking trend was observed in that the larger motifs were clearly more stable by all measured. To objectively assess this, cohort means were compared for statistical significance. In all cases, 1- and 2-STR motifs were significantly destabilized with regard to wild-type D20:24; for all but the PK_50_ measure they were also destabilized with regard to 3- and 4-STR cohorts. The situation with regard to 3- and 4-STR cohorts is more complex, since they were more similar to D20:24 and also since these cohorts also had a much greater variance in all parameters. This resulted in no significance on a cohort basis between 3- and 4-STR targets and D20:24 except for f_α_.

Overall, this shows that in this region at least, a larger 3-STR minimum size is a necessary, but not sufficient condition for wild-type like stability. This also begs the question – why not expand the size for all edits to at least 3-STR? This is impossible since the dystrophin rod is divided and delimited by various disordered “hinge” regions that break up the continuous STR structure and prevent structural interaction. These hinges occur before D1, hinge 1; between D3 and D4, hinge 2; between D19 and D20, hinge 3, and after D24, hinge 4; with a non-classical hinge-like region also occurring In the D20 to D24 region of this study we are thus bounded by H3 and H4. It remains to be seen if this size correlation occurs in other regions of the rod, especially the large region bounded by hinges 2 and 3, D4 to D18.

### Alternative Exon Skip Repairs for Same DMD Deletion

Within our cohort of 11 edits, 6 of them are of special interest, since they are end products of alterative exon skip repairs of a single DMD type defect. These occur in overlapping pairs, linked to 4 specific DMD out-of-frame defects listed in Table 4. For patients with these defects, other good or bad edits are largely irrelevant, as is the wild type protein. These patients only have access to these two edited choices so far as exon skipping therapy is concerned. Indeed, with the recent approvals of golodirsen and viltolarsen (12, 13), this is now a clinical reality for some patients, albeit not for the edits in this study.

**Table 4.** Alternative repairs in this study. The DMD type, out-of-frame genetic defect can have their reading frames corrected by skipping a flanking exon int two ways, resulting in two possible exon edited proteins.

| DMD defect | Exon skipped | Exon edit |
| --- | --- | --- |
| $\Delta e53-57$ | exon 52 | $\Delta e52-57$ |
| | exon 58 | $\Delta e53-58$ |
| $\Delta e54-58$ | exon 53 | |
| | exon 59 | $\Delta e54-59$ |
| $\Delta e55$ | exon 54 | $\Delta e54-55$ |
| | exon 56 | $\Delta e55-56$ |
| $\Delta e56$ | exon 55 | |
| | exon 57 | $\Delta e56-57$ |

With these alternative skip targets, there are two sets: a group of three, Δe54-55, Δe55-56 and Δe56-57 applicable to Δe56 and Δe57 DMD defects; and a second group of three Δe52-57, Δe53-58 and Δe54-59, applicable to Δe53-57 and Δe54-58 defects (see Table 4). In the first case, Δe55-56 is clearly a superior edit with respect to fidelity with wild-type, undamaged D20-24, Figure 7. It has high helicity, 93%, similar to Δe56-57 and only a bit shy of the wild type value of 98%. With regard to tertiary structure stability, it is very stable (in fact *more* stable than D2024) whereas both of the other alternatives are significantly less stable; and similarly, it has a high, wildtype-like PK_50_ value indicating few if any disordered regions, similar to wild type rods, whereas the other two have significantly lower PK_50_ values is indicating more disrupted structure. By all three measures Δe55-56 is more stable and comparable to wild type. While the alternatives, Δe54-55 and Δe56-57, are destabilized, and more perturbed.

**Figure 7.**
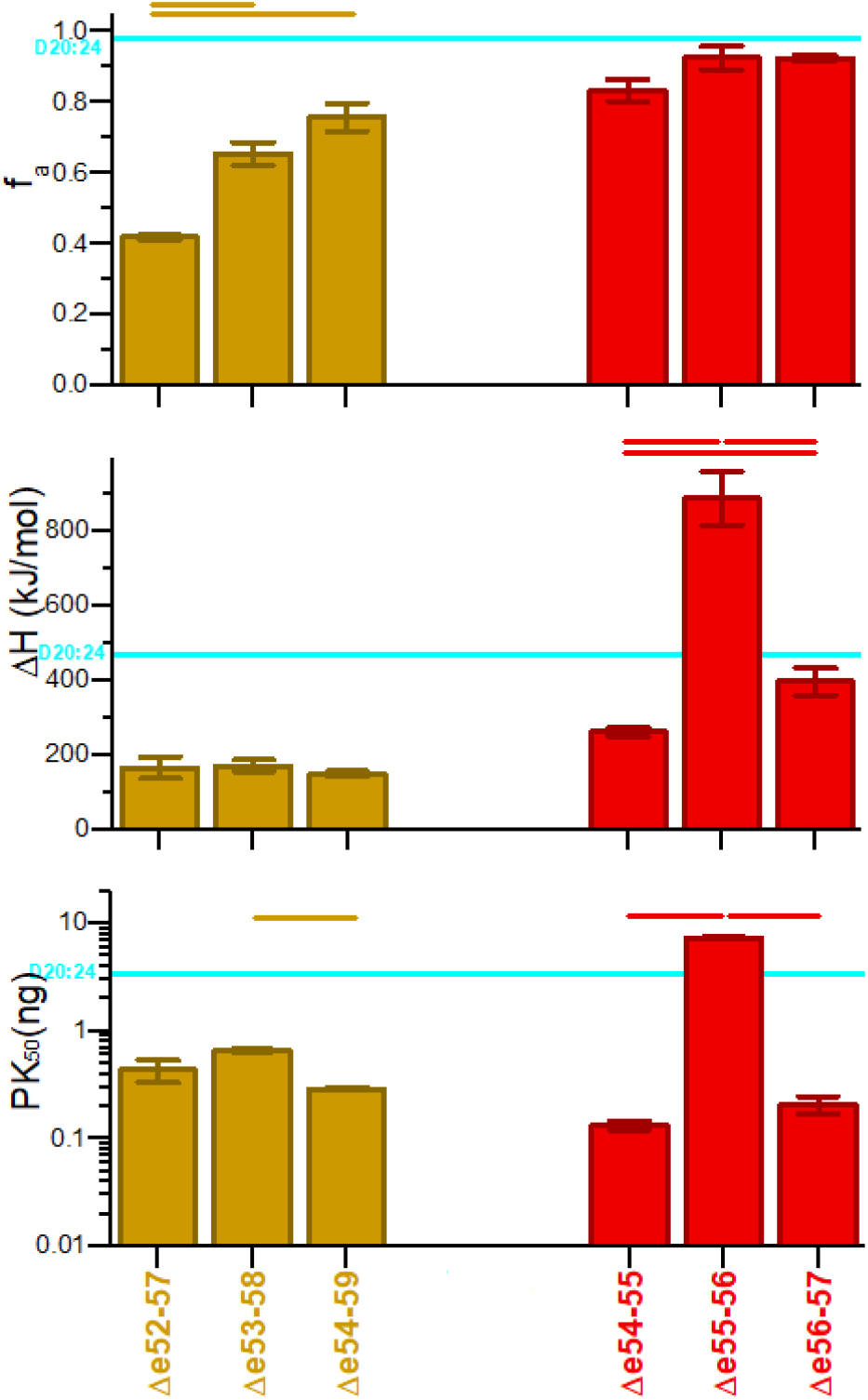
Alternative exon skip targets. Our cohort contains two groups of alternative-repair edits, and we compare them here with regard to secondary structure, fa; tertiary structure stability, DH; and presence of disordered region PK_50_. For reference, the value for unperturbed D20:24 is shown as a blue line. In one group (red, right) a clearly less perturbative edit is evident: De55-56 which is more stable and wild type like with regard to both measures. In fact, by some measures, DH and PK_50_, it is more stable than wild type. In the other set (gold, left) there is no clear best choice and all edits are significantly destabilized with respect to wild type values. Statistically significant differences, P<0-.005, are indicated by bars above.

In the second case, however, there is no clear choice. All of them have helicity values well below wild type, with Δe54-59 the best at 76%, compared to 98% for wildtype; and they all have destabilized tertiary structure with ΔH value below 171 kJ/mol (D2024_WT_: 467 kJ/mol). Indeed, in two cases, Δe52-57 and Δe54-59, two distinct low enthalpy transitions are observed (see Figure 3) indicating that there are two separate non-cooperative folding units, separated by some disordered linker region – presumably at the edit site. This is supported by very low PK_50_ values for these two targets in particular, but also Δe53-58 - all of which are < 0.7 ng, well below wild type values (3.4 ng). In this set of alternatives, unfortunately, there is really no good choice for an edit that can mimic damaged dystrophin.

Overall, this shows that in some cases, alternative repairs can have very different properties, with in some cases one choice being much more wildtype-like than the other. It is yet to be established whether this will have clinical ramifications in these specific cases; however other studies have suggested a significant link between perturbed STR structure and clinical progression (20, 41). Equally significant, the fact that in some cases, no clearly superior wild-type choice emerges, and alternative edit strategies may be necessary. This heterogeneity has been seen in other studies, with in some cases a clear differences in alternatives seen (22, 31), and in other cases no clearly superior best edit emerging (29). However, this is the first study to produce both results in one large cohort, and confirms that this heterogeneity may simply be a property of the specific region being edited.

### Hybrid STRs vs. Non-hybrid (Junction) STRs

An additional factor we consider is the status of an edit as “hybrid” or “junction” type edit. There are approximately two exons per STR, so that deletions removing an integer number of STRs generally remove an even number of exons and are referred to as in-phase (note not in-frame which refers to the reading frame). On the other hand, deletion removing an odd number of exons leave behind a presumably disordered fractional STR, these are called out-of-phase. In-phase edits have been shown to result in significantly milder clinical phenotypes in BMD patients than out-of-phase (20). All the edits studied here are in-phase, which is simply an outcome dictated by specific exon reading frames in the D20-24 region. However, in-phase edits may be further categorized. Since there are ~2 exons per STR, exon boundaries can be classified as either “middle” or “junction”. When an in-phase edit joins a junction to a junction boundary the result is an edit approximately at an STR junction, and the neat excision of intervening STRs. However, when an edit joins a middle to a middle, this fuses the N-terminal region of one STR to the C-terminal past of another STR resulting in a hybrid STR. These hybrid STRs have been shown to be viable, but are often destabilized relative to junction type edits.

In this study, due to the nature of the exons in the D20-24 region, all but one edit resulted in hybrid STRs; this can be seen in Figure S1 where all edits are presented in a fashion that shows the edit site in relation or modelled STR structure. The sole junction type edit was Δe55-56. It was indeed more stable that all other, hybrid type edits. This was statistically significant every case, with the sole exception of helicity where significance was seen only in 6 of 10 comparisons (see Figure 8). This once again demonstrates that junction type edits appear to be more likely to produce stable structure than hybrid edits. However, we do note they some hybrid edits are much better than others, and approach native rod stability.

**Figure 8.**
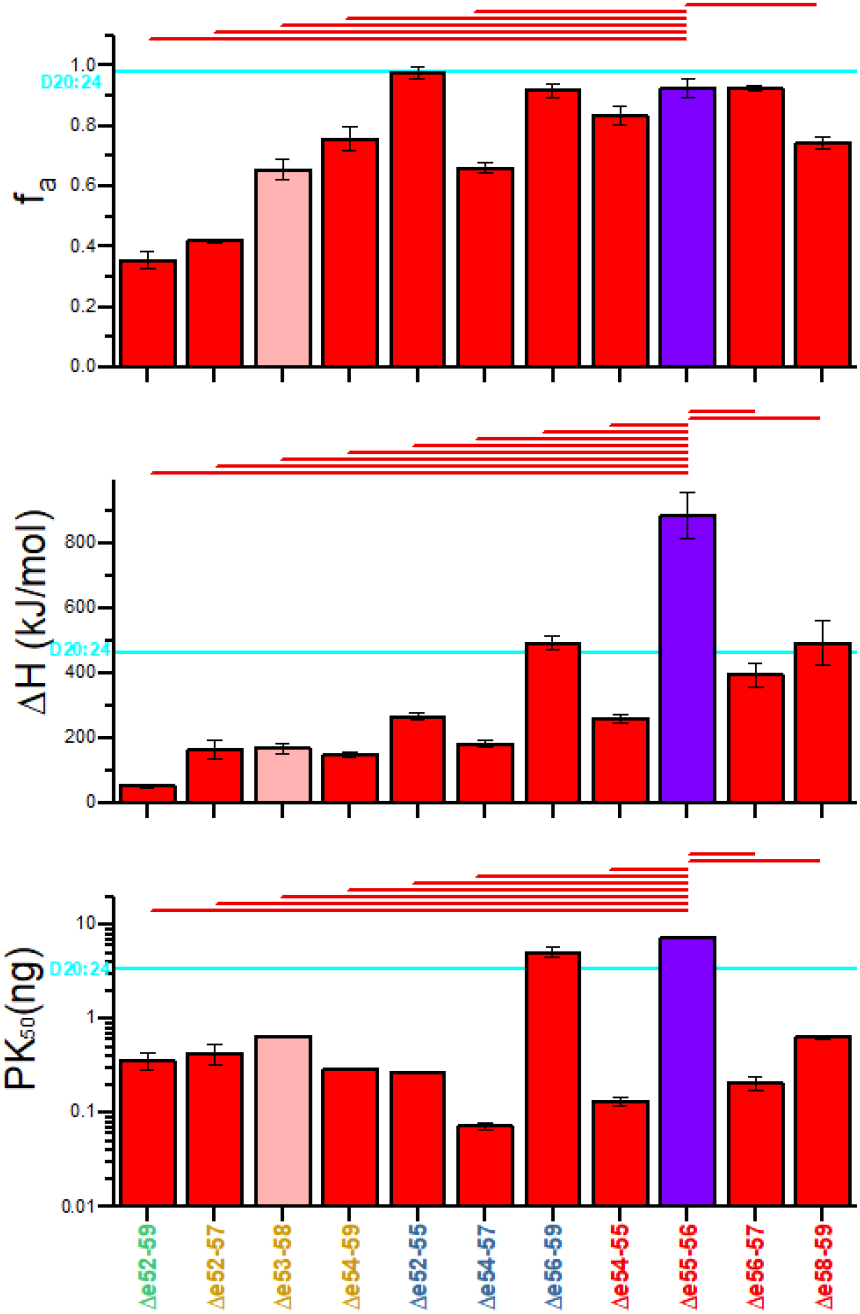
Junction vs hybrid STR in-phase edits. In the study region there is only one true junction edit, Δe55-56, blue; hybrid edits are in red. The Δe53-58 edit, which fuses compatible “STR end” exon boundaries might at first thought be considered a junction edit, but due to heterogeneity on the length of STRs which have only limited homology, the edit site is located well within the STR structure in modelled structures. (See figure S1 for structures and S2 for exon boundaries). We thus classify this as a hybrid edit, but highlight this distinction by lighter shading. We see that the junction edit is less perturbed and more wildtype like (D20:24, blue line) than most hybrid edits. Statistical significance (P<0.05) is shown by bars above.

### Proteolytic sensitivity and D23 loop

One metric, proteolytic sensitivity, provided some interesting results, since two edits (Δe55-56: 7.40ng, Δe56-59: 5.14ng) displayed significantly increased resistance to proteolysis compared to native protein (D20:24: 3.42ng). Both of these were also thermodynamically more stable than wildtype; significantly so (P<0.005) for Δe55-56, but not significantly so for Δe56-59 (495 vs 476 kJ/mol, P=0.06). This is very unusual, since in all exon edits so far studied, none were stabilized: not in the exon 43-45 region, not in the exon 51 region (29), and not in larger deletion from exon 44-56 region (16, 22). To examine this, we examined Robetta models of all edits with at least 3 STRs for disordered loop regions, which are known to correlate with proteolytic susceptibility. This is really the whole point of the proteolysis assay: to detect such locally perturbed region, even when the rest of the molecule folded well and might dominate the thermodynamic analysis. We found that all motifs with low PK_50_ values exhibited such loop region but none of those with high PK_50_ values did (Figure 9). Furthermore, we also determined that these very same motifs also display thermodynamic stability differences in the same clustering.

**Figure 9.**
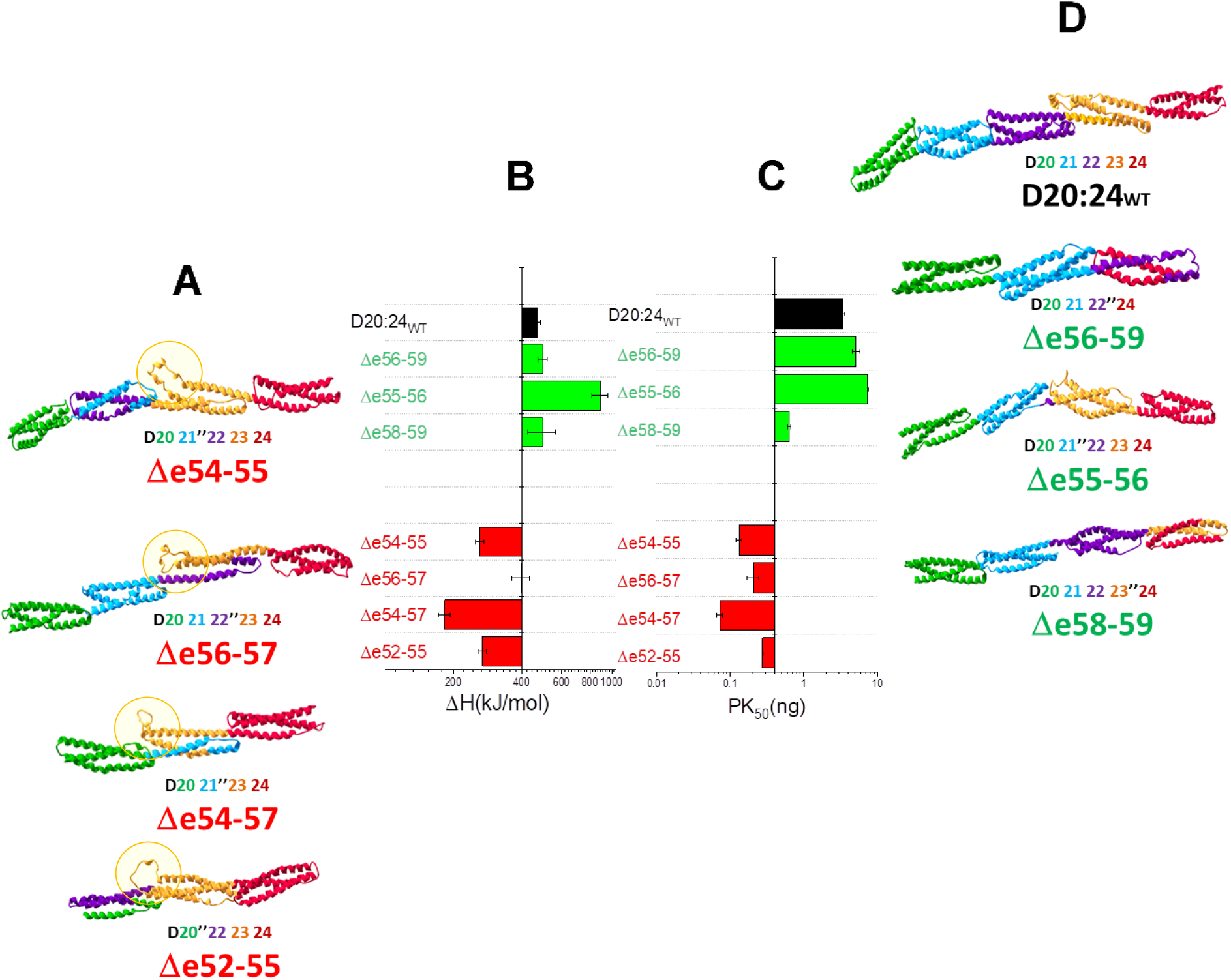
Robetta models targets with more than 3 STRs, as well as D20:24 compared to PK_50_ and ΔH. These are grouped into low PK_50_ cohort, **A**, and a high PK_50_ cohort, **D**. In the middle are shown unfolding enthalpy ΔH, **B**, and PK_50_ **C** bar graphs. The low stability cohort in all cases contained a disordered loop, yellow circle, in the D23 region, yellow. This loop occurs in a similar position, in all cases: in the turn between the second, B, and third, C helix of the STR containing the D23 sequence, which places it at the end of this STR that interacts with the previous STR. In two cases, Δe56-57 and Δe54-57, a hybrid STR forms in the model so it is perhaps not surprising that part of the D23 sequence is not fully integrated. In the two other cases, Δe54-55 and Δe52-55, the edit site is distal to D23, but creates a hybrid STR immediately prior to D23, which appears to not interact with this region n D23 as productively as native D22 appears to, in the wild type D20-24 rod.

### Conclusions

In this study we have examined 11 possible exon edits in the D20-24 regions. This has produced further evidence that different exon edits impact protein structure to different extents. This is not overly surprising, since several studies have shown this in a more limited fashion; however, this study was the first to examine a systematic panel of all possible edits in a given region. We also for the first time demonstrate two exon edits of stability comparable to, and by some measures greater than, the wild type protein rod. These are the Δe55-56 and Δe56-59 edits, which are the least perturbative exon edits yet identified. They provide hope that fully functional synthetic edited dystrophins might be possible.

We have also shed light on some factors that may contribute to more functional edits. We show that hybrid type edits are in general less stable than junction type edits, but also clearly demonstrate that hybrid edits are very heterogeneous and that some approach wildtype and junction type edits in stability. It seems likely that highly specific atomic level interactions are necessary to form a hybrid STR out of two fractional STRs, and that hybrid STR stability will be very sensitive to the specific pairing of STR fragments. Other work in the exon 45-47region (42) has suggested that the dystrophin rod can be very sensitive to helix-helix interactions and that in frame deletions that perturb these can be generally quite disruptive, resulting in loss not only of structural stability, but also binding interaction, and this one case an important binding interaction with nNOS. In contrast, junction type edits do not perturb the core of any STR structure, so they may be better tolerated. However, for many patients, exon skipping only leads to hybrid type exon edits.

In this study we have further examined alternative repair pairs, which occur when, for some types of DMD defects, exon skipping can be applied in more than one fashion. These are especially crucial as additional exon skipping therapies are developed. In addition to eteplirsen which skips exon 51, golodirsen (13) and viltolarsen (12), both targeting exon 53, have recently been approved in the US. Thus, for some patients (those with an exon 52 deletion) this is now a clinical reality. Reagents targeting other exons are in late stage clinical trials, so this is likely to become a larger issue. Here we find evidence that in some cases, (as previously found) a clearly superior choice in terms of protein level perturbations exists; whereas in other cases (also as previously found) there appears to be no good choice. In the end what we have shown is that the dystrophin STR rod is highly complex, and empirical study in necessary to determine the properties of exon edits, many of which may have clinical ramifications.

We further show, at least in the D20-24 regions, that longer tandem STR units are more stable than shorter ones, with a length of ~ 3STR necessary but not sufficient to approach wildtype in stability. This revives a long standing conceptual debate in STR containing rods: should they be viewed as independent, triple α-helix bundles (43), or should he be viewed as a complex fiber, composed of a nested series of long α-helices in one direction (i.e. forward) alternating with shorter α-helix in the other direction (i.e. backwards) (26). The former view, implicit in how STR are described and annotated in many data bases, suggests they are independent units and can be shuffled about easily; the latter nested view suggest that adjacent STRs will be structurally integrated and cooperative in folding which may make editing more complex. Indeed, a full thermodynamic scan of the dystrophin STR rod (23) suggests that both models are applicable in different region of the overall rod; but that the nested, interacting model applies in the D20-24 region. As such, in this region it is perhaps not surprising that we have found that longer rods with such inter-STR interactions are less perturbed. This however has consequences when considering optimal exon-edit therapy, or in designing synthetic edited dystrophins.

## Supplemental Figures

**Figure S1.**
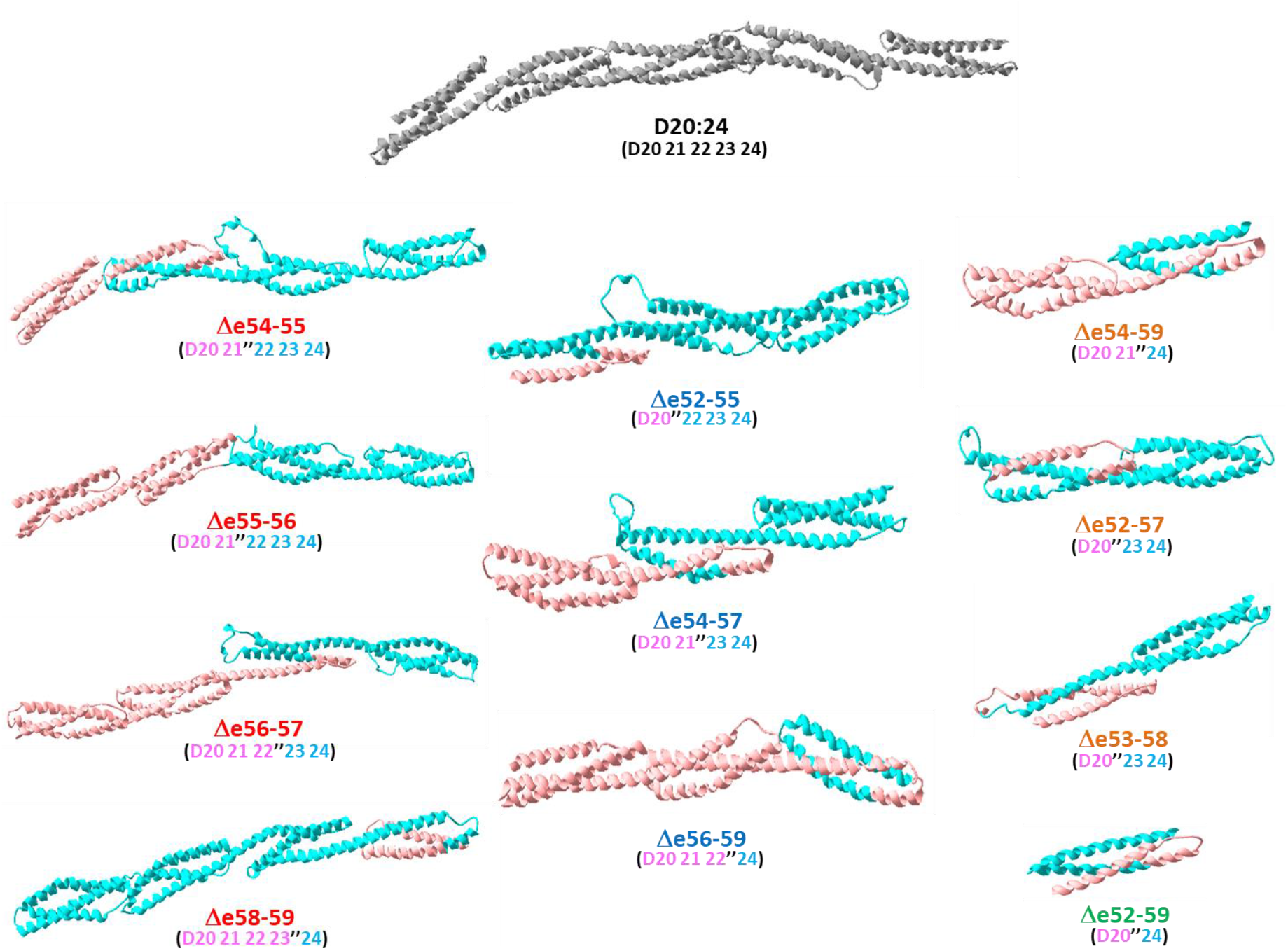
Target rod models showing edit site in relation to STR structure. Target proteins were modelled using the Robetta protein modelling server. At the top, the 5-STR D20:24 unedited rod is shown in grey. Below, the eleven target exon edited proteins are shown with the N-terminal region before the edit site in pink, and the C-terminal region after the site in blue. Also shown below the name of each edit is the STR structure in the same coloring scheme. Partial STRs are denoted as X’ (N-terminal region) or ‘Y (C-terminal fraction). Thus, hybrid STRs become X”Y. We can see that only Δe55-56 has the exon edit junction at an STR junction, between the 2^nd^ and 3rt STR there. All others have edit junctions internal to STRs, so creating a hybrid STR at the edit site.

**Figure S2.**
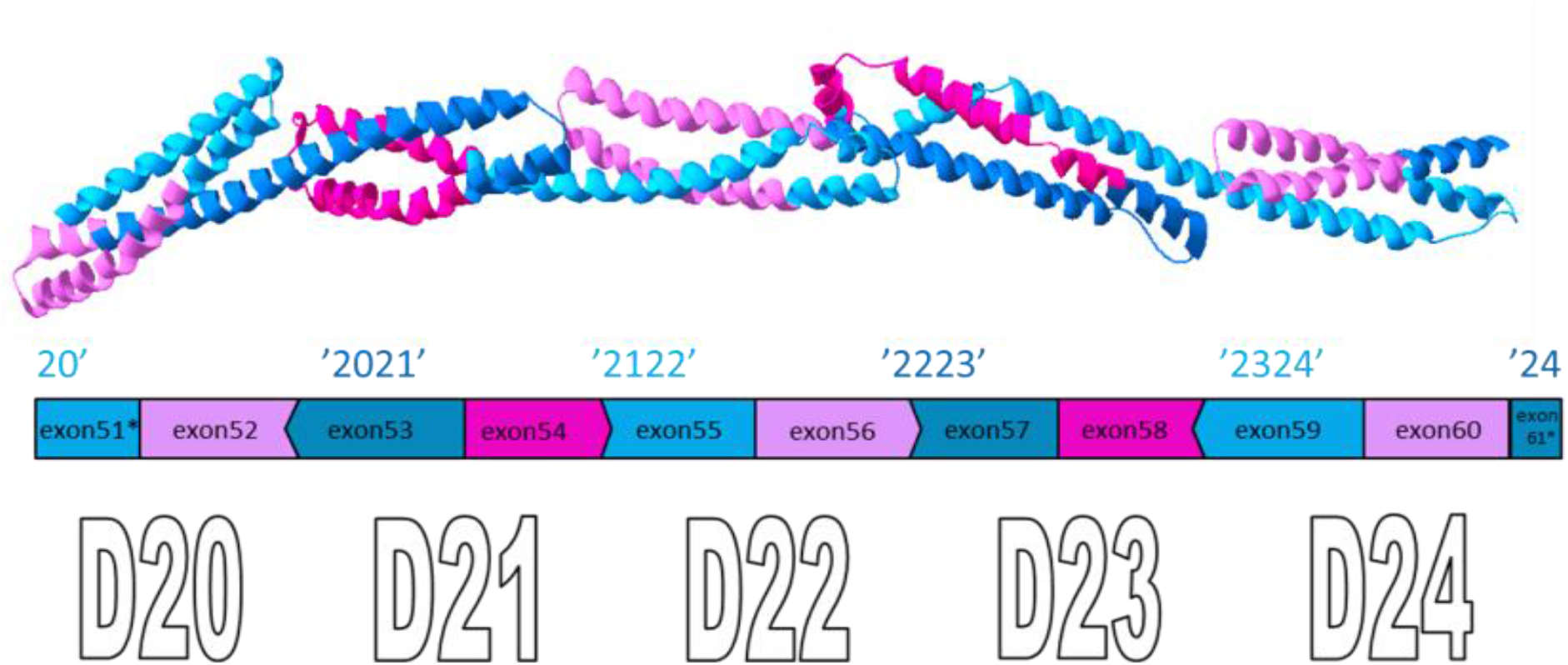
STR structure in relation to exon structure. The modelled D20:24 native rod consists of five STRs, and is presented with the exons colored alternately magenta and blue (the intensity of each is modulated for clarity, where they overlap in the 3-dimnesional modelled structure). We see that the model is consistent with a unified rod, where the last helix of one STR propagates directly into the first STR of the next. Exon reading frame is shown by the shape the vertical edges for the exons: **|** is frame 0 (i.e. at codon boundary); **>** is frame 1 (i.e. between 1^st^ and 2^nd^ base of a codon). **<** is −1 frame (i.e. between 2^nd^ and 3^rd^ base of a codon). Viable edits need to match up compatible edges to ensure proper reading frame.

**Figure S3.**
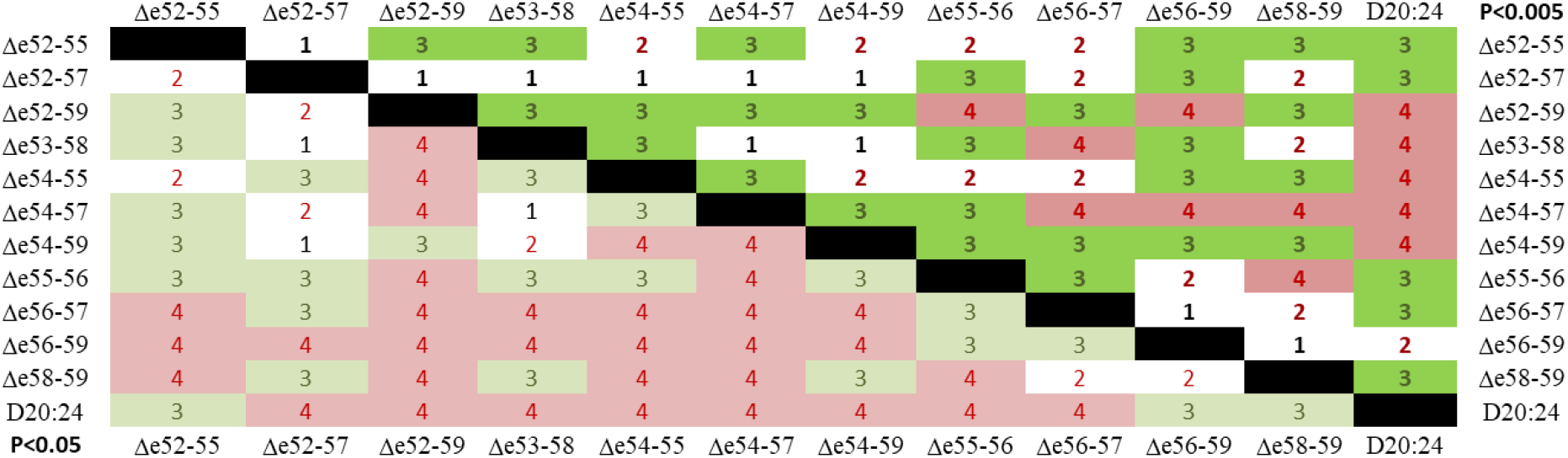
Significance of Differences in Edited Rod Properties. The number of comparisons of four protein property parameters (f_a_, Tm, ΔH, PK_50_) among all 12 rods (D20:24 wild-type plus 11 exon-edit targets) that are statistically significant for all pair-wise comparisons is presented. In the upper right triangle the standard was *highly significant*, P<0.005; and in the lower left the standard was *significant* P<0.05. Of the 66 pairwise comparisons, 50% of them show significance at P<0.05 in all four measured properties, and 85% in at least three. For highly statistically significant differences. P<0.005, 18% achieved this in four measurements, and 65% in at least three methods. This emphasizes that the nature of the edit site has a large impact on protein properties.

